# Unveiling Hidden Connections in Omics Data *via* pyPARAGON: an Integrative Hybrid Approach for Disease Network Construction

**DOI:** 10.1101/2023.07.13.547583

**Authors:** M. Kaan Arici, Nurcan Tuncbag

## Abstract

Omics technologies are powerful tools for detecting dysregulated and altered signaling components in various contexts, encompassing disease states, patients, and drug-perturbations. Network inference or reconstruction algorithms play an integral role in the successful analysis and identification of causal relationships between omics hits. However, accurate representation of signaling networks and identification of context-specific interactions within sparse omics datasets in complex interactomes pose significant challenges in integrative approaches. To address these challenges, we present pyPARAGON (PAgeRAnk-flux on Graphlet-guided network for multi-Omic data integratioN), a novel tool that combines network propagation with graphlets. By leveraging network motifs instead of pairwise connections among proteins, pyPARAGON offers improved accuracy and reduces the inclusion of nonspecific interactions in signaling networks. Through comprehensive evaluations on benchmark cancer signaling pathways, we demonstrate that pyPARAGON outperforms state-of-the-art approaches in node propagation and edge inference. Furthermore, pyPARAGON exhibits promising performance in discovering cancer driver networks. Notably, we demonstrate its utility in network-based stratification of patient tumors by integrating phosphoproteomic data from 105 breast cancer tumors with the interactome, leading to the discovery of tumor-specific signaling pathways. Overall, the development and evaluation of pyPARAGON significantly contributes to the field as an effective tool for the analysis and integration of multi-omic data in the context of signaling networks. pyPARAGON is available at https://github.com/metunetlab/pyPARAGON.

## Introduction

Omics technologies provide a multidimensional view of the cell’s functional mechanism, context-specific alterations, such as in diseases or drug perturbations, and biological processes ^1,2^. As the omics data accumulate, integrating them accurately and translating them into interpretable knowledge remains challenging due to data sparsity, missing data points, and computational complexity ^3–5^. Omic hits are sparsely connected in a reference interactome and carry noise from high-throughput outcomes ^6,7^.

Recent methods utilizing learning- and network-based algorithms are on the rise to overcome these challenges and decode causal relations between omic entities ^8–11^. Learning based methods efficiently integrate multi-omic data to extract interpretable annotations such as pathways, reactions, and processes ^12–14^. Also, network-based algorithms, including shortest paths ^15^, Steiner trees/forests ^16,17^, and random walk ^18,19^ have been frequently used to construct specific networks by propagating omic hits ^20,21^. Network-based methods can uncover the most relevant interactions between a given set of proteins/genes by either inferring from a reference protein-protein interaction (PPI) network or reconstructing them ^1,22,23^. These methods eventually obtain a network model which may represent the alterations in disease models or drug treatments with the help of topological and statistical features ^24–29^. The benefit of using global and local network features (e.g. degree distribution, clustering coefficients) for propagation or inference ^30,31^ is limited when this type of sparse data is elaborated ^32,33^. Therefore, the frequency of motifs (repeating subgraphs) can be more explanatory for deciphering complex cellular networks ^34,35^. Small connected, non-isomorphic subgraphs, called *graphlets*, are over-represented in the reference interactome and associated with specific functions ^35,36^. Graphlet statistics solve several complex problems in this context, such as the comparison of biological networks, delineating the functional organization of networks, discovering functionally related genes, regulatory interactions, and parameter tuning for network-based approaches ^12,32,33,37–40^. Another challenge is the presence of highly connected and multifunctional proteins, particularly hub proteins, which can bring nonspecific interactions to the resulting network models because of the small-world property of reference interactomes. Therefore, using network motifs, graphlets, or revealing modules can improve the context-specific aspects of the models ^1,25,41^.

In this study, we hypothesize that the utilization of network motifs, in lieu of pairwise connections among proteins, may provide a more accurate representation of signaling networks and mitigate the inclusion of nonspecific interactions. Therefore, we present pyPARAGON (PAgeRAnk-flux on Graphlet-guided network for multi-Omic data integratioN) as a new tool that combines network propagation with graphlets to construct context-specific networks. We demonstrated that graphlets reduce the dimensionality of complex reference interactomes by filtering out non-specific and highly connected proteins and their interactions. pyPARAGON performed better in node propagation and edge inference than selected state-of-the-art approaches, including Omics Integrator 2 (OI2) and PathLinker (PL) on the benchmark set of cancer signaling pathways. pyPARAGON also showed promising results in the discovery of cancer driver networks. Finally, we showed the network-based stratification of patient tumors as a use case of pyPARAGON and found tumor-specific signaling pathways when phosphoproteomic data from 105 breast cancer tumors were integrated with the interactome.

## Results

### Overview of pyPARAGON as a hybrid network inference framework

Recent studies have shown that when integrating different types of biological data (such as genomic, proteomic, and transcriptomic data) to reconstruct signaling networks, using hybrid approaches can be more effective than relying on a single method alone ^16^. The accuracy of reconstructed networks is also highly dependent on the quality of the reference interactome ^42,43^. However, there are tradeoffs involved in increasing the number of interactions in the reference interactome. On the one hand, including interactions with low confidence scores may lead to the identification of false positive proteins and interactions. On the other hand, highly connected proteins (i.e., hubs) may dominate the final network and obscure context-specific relationships between proteins or genes. Graphlets are small, connected subgraphs with a specific pattern of edges and are similar to network motifs in that they represent recurring patterns ^35,36^. Here, in pyPARAGON, we present a new hybrid approach that combines graphlets with network propagation *via* the personalized PageRank algorithm, followed by interaction selection based on edge flux calculation, to address these challenges in network modeling. pyPARAGON has three steps (**Figure 1A**): i. Graphlet-guided network (GGN) construction; ii. Propagation and edge scoring *via* the Personalized PageRank (PPR) algorithm and flux calculation; iii. Subnetwork construction with highly scored edges on GGN. pyPARAGON takes a list of seed genes/proteins (initial nodes) as input, which are specific to the biological context of interest. Seeds can be obtained from but not limited to omics experiments, drug perturbation analysis, or disease-associated proteins. Graphlets help in reducing the load of possible false positives, while network propagation via PPR scores all other proteins in the reference interactome by orienting them from the given seed list. This hybrid approach, on one hand, trims the reference interactome to the most relevant interactions by constructing a GGN, on the other hand, it quantifies the importance of other proteins based on a given seed list.

**Figure 1.**
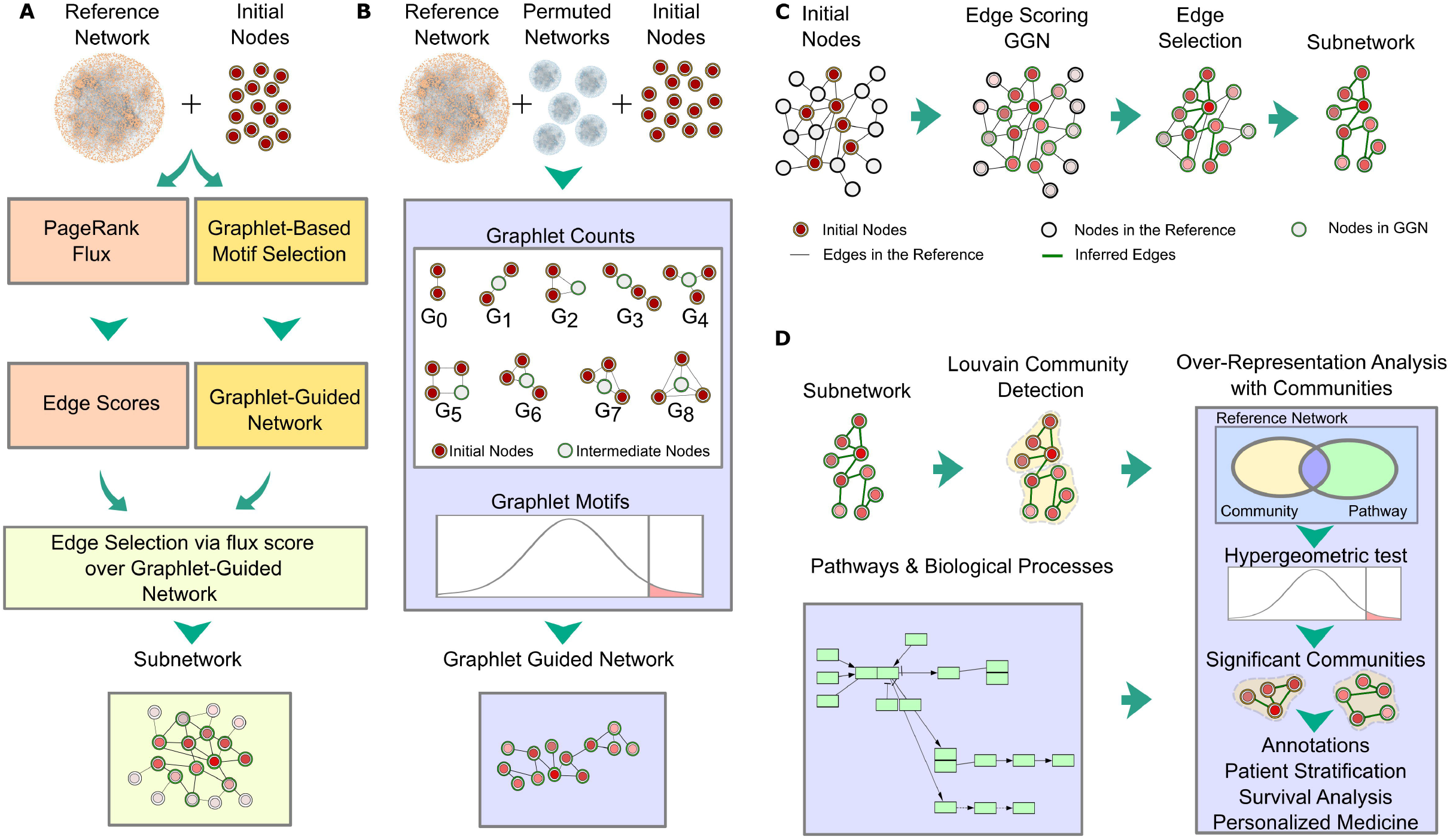
The overview of pyPARAGON **a.** pyPARAGON has three steps: i. GGN construction (light red boxes); ii. Edge scoring with personalized PageRank flux calculation (yellow boxes); iii. Subnetwork inference using edge scores and GGN (green boxes). **b.** We investigated nine non-isomorphic graphlets (G_0_-G_8_) composed of 2, 3, and 4 nodes for GGN. Except for G_0_, each graphlet covers at least two seed nodes (red circles) and one intermediate node (white circles) that connects the seeds in the center of the orbit. To find significant motifs, we screened the presence of each graphlet in 100 randomly generated reference interactomes using the same seed nodes. We tested the significance of each graphlet’s presence in real interactomes compared to random interactomes with the z-test (p < 0.05, z-score > 1.65). Significant graphlets were merged to construct GGN. **c.** By random walking from weighted initial nodes in the reference network, the Personalized PageRank algorithm assigns a weight to each node. Computed edge fluxes were used as the edge scores in the reference interactome. High scoring edges in GGN formed the final subnetwork. **d.** The network analysis module of pyPARAGON employs Louvain community detection methods, based on network topology, to divide the inferred network into functional units. Significant biological processes and pathways in each community were found by hypergeometric test.

In general, state-of-the-art methods use an immediate edge between two nodes in the reference network and node-based features (e.g. degree, betweenness, closeness, and eigenvector centralities). GGN construction step of pyPARAGON goes beyond this and follows an unsupervised approach to identify a core region in the reference interactome by combining significantly frequent graphlets composed of 2-, 3-, and 4-nodes (**Figure 1B**). In omics-based network construction, direct connections between the genes/proteins of interest are often sparse, and intermediate nodes are required to connect them and form a coherent network structure. Thus, we constrained that graphlets having more than two nodes may have an intermediate node. Intermediate nodes are the ones that have the highest connections to the seed nodes in the corresponding graphlet.

In addition to GGN construction, the personalized PageRank algorithm propagates signals from seed nodes across the reference interactome. Node weights after propagation together with their degrees and edge confidence scores are combined in a single function to calculate edge fluxes (see Methods) ^44^. In this function, the degree component penalizes highly connected proteins that are non-specifically present in the resulting subnetworks. Finally, we map edges with high flux to GGN to obtain a context-specific network (**Figure 1C**). pyPARAGON also uncovers communities functioning in specific biological processes or pathways (**Figure 1D**). Based on network topology, the Louvain community detection method divides inferred subnetworks into small modules ^45^. Then, using a hypergeometric test, pyPARAGON discovers context-specific annotations ^46^. In this way, we reveal not only hidden connections between initial nodes but also significant context-specific pathways.

### Network Trimming via Graphlets Improves the Network Inference

Benchmark datasets for assessing the performance of network methods are usually curated biological pathways. In general, the performance of the methods is evaluated based on topological features, coverage of predicted nodes, and edges. We used NetPath ^47^ as the benchmark dataset to reconstruct curated cancer signaling pathways and assess the performance of pyPARAGON. As a result of screening all graphlets across the reference interactomes, we found ***G_2_***, ***G_5_***, ***G_6_***, ***G_7_***, and ***G_8_*** to be the most frequent graphlets (**Supplementary Figure 1A**). The frequency of direct interactions between input nodes (represented with ***G_0_***) is not significant in the reference interactome; however, the presence of these direct interactions in a graphlet with at least three nodes is significant. For example, the direct interaction of seed nodes in ***G_2_***is more significant in the presence of an intermediate node interacting with ***G_0_***. As to our observation, significant graphlets having at least one intermediate node to connect seeds provide more precision compared to including direct interactions between two seeds (i.e. ***G_0_***) in GGN.

Each available interactome has a specific evaluation and scoring scheme to integrate PPIs from different resources ^42^. In this study, we used ConsensusPathDB, HIPPIE v2.2, and HIPPIE v2.3 which have different topological features (**Supplementary Table 1**). The constructed GGN by pyPARAGON is a subnetwork of the reference interactome. When we compared the original interactomes and trimmed interactome via GGN construction separately, we observed that the similarity across them significantly increases when GGNs are used (**Supplementary Figure 1B**). Another advantage of GGN construction is attenuating the dominance of the highly connected nodes. Notably, highly connected proteins have numerous functions in the cellular processes known from prior knowledge ^48^. We identified 1812 highly connected proteins in HIPPIE v2.3 (nodes having more than 200 interactions). With the help of graphlets, GGN construction step in pyPARAGON successfully trimmed many interactions and decreased the number of highly connected nodes significantly (**Figure 2A**). Protein interactomes are scale-free networks intrinsically following a power law (**Supplementary Figure 1C**) ^49^. The resulting GGNs in pyPARAGON preserve this property of scale-free networks.

**Figure 2.**
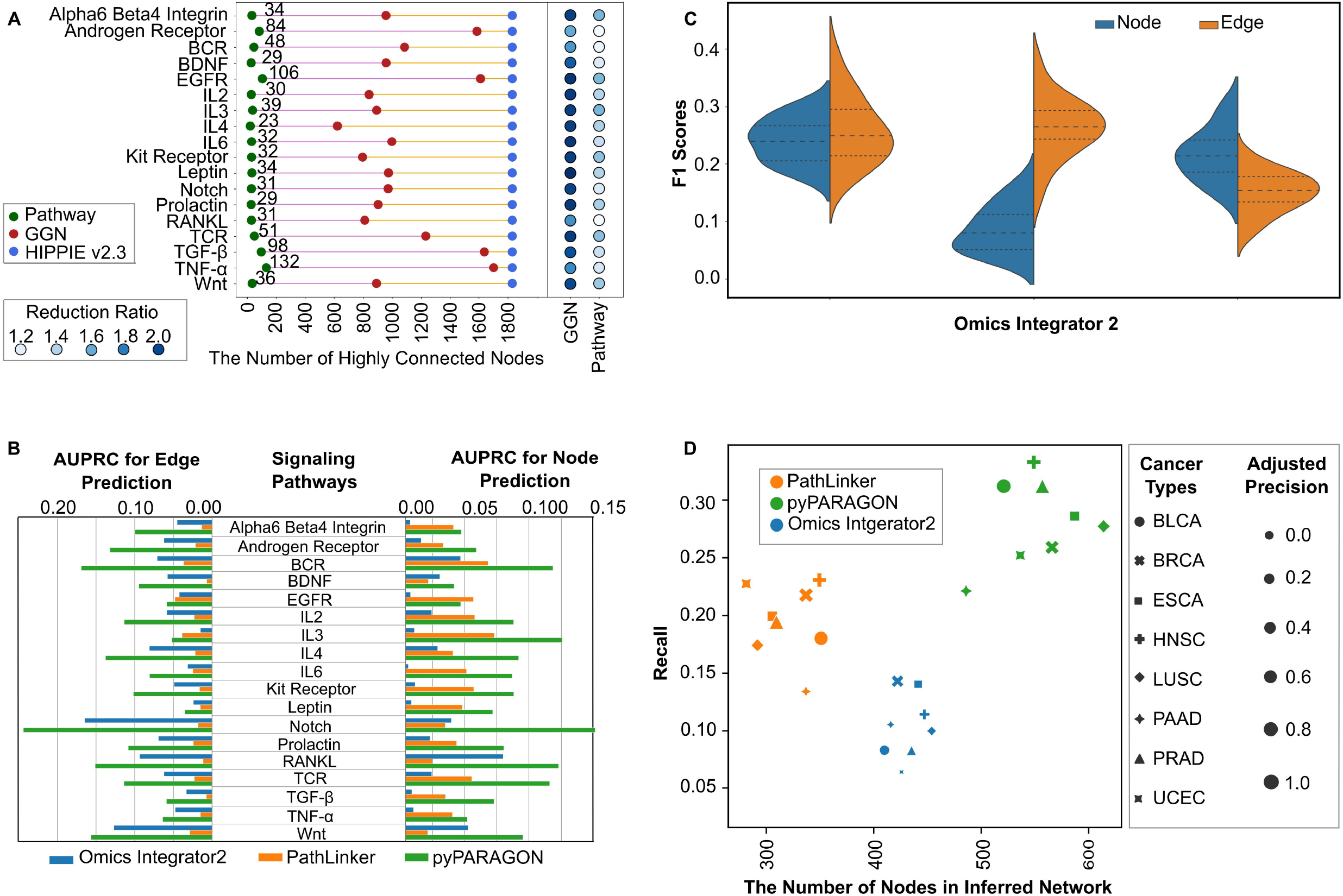
Graphlet-guided network trims reference interactome by removing some highly connected nodes and their non-specific interactions. **a.** Highly connected proteins are defined as the ones having more than 200 interactions in HIPPIE interactome (blue dots). Presence of these nodes in GNNs and NetPath pathways are shown for each signaling pathway (red and green dots, respectively). In the reference interactome, 1812 highly connected nodes are present. GGN selects a subset of these nodes that are highly specific to the pathways. The change in node degrees of remaining highly connected proteins in GGN was calculated as the reduction ratio and shown with a blue color scale. Highly connected nodes in the reference interactome that are present in pathways are included during reconstruction with a low reduction ratio in GGN while the rest have a higher reduction ratio. **b.** AUPRC of each tool (blue= OI2, orange=PL, and green=pyPARAGON) in each pathway reconstruction is shown in barplot. In all signaling pathways, pyPARAGON performed better than others in both node and edge predictions. **c.** Distribution of F1-scores for each tool across 18 pathways is shown for node (blue) and edge (orange) predictions **d.** We inferred cancer-specific networks for eight distinct cancer types by using the most commonly mutated genes as initial node sets. Marker size represents precision, while recall and network sizes are shown on the x-axis and y-axis. The recall score represents the ratio of correctly predicted cancer driver genes in cancer-specific networks to total number of drivers. pyPARAGON achieved better recall scores for each cancer type without having a decrease in precision scores.

We conducted a comparative evaluation of pyPARAGON against PathLinker (PL) and Omics Integrator 2 (OI2) to infer curated signaling pathways in NetPath. Since there is no definitive benchmark or ground truth for assessing tool performance, we relied on propagated nodes and predicted edges as evaluation criteria. We measured performance using the area under the precision-recall (AUCPR) curve to demonstrate how well each pathway’s nodes and edges were recovered in the predicted networks. Our analysis showed that pyPARAGON outperformed PL and OI2 at both the node and edge levels for inferring signaling pathways in all pathways of NetPath (**Figure 2B**).

Performance comparison of pyPARAGON with others was done in two directions: i. node propagation, ii. edge inference. We used the F1 score to compare them because it simultaneously represents precision and recall in one metric. The overall results show that pyPARAGON and PL are better at propagation, while pyPARAGON and OI2 are better at network inference (**Figure 2C**). Highly connected reference networks decreased the propagation ability of OI2 while providing more robust interactions than PL. On the other hand, PL propagated the seed node set more robustly due to the consideration of multiple short paths but introduced many false positive interactions. Many seed nodes have a tendency to be connected by hub nodes as shortcuts due to biological networks being scale-free. Thus, the shortest path and random walk-based approaches may lead to including false positive interactions ^20,50^. However, penalizing highly connected nodes, e.g. calculation of PageRank flux normalized the score in pyPARAGON or degree-dependent negative prizing in OI2, reduces false positive edges and improves F1-score in edge prediction

Next, we used pyPARAGON to predict cancer driver pathways and compared its performance with other selected tools. Here, we labeled 300 genes with the most prevalent mutations in eight cancer types as seed nodes. Known driver genes in IntOGen were used as an independent test set for performance evaluation based on their presence in the reconstructed networks. Because we use 5-fold cross-validation, for each fold we filtered out the common proteins between the seed list and known drivers and then reconstructed cancer type-specific networks with pyPARAGON, PL and OI2.

Our results show that the reconstructed network by pyPARAGON covers more known driver genes compared to other methods and achieves higher recall and precision compared to other methods in all cancer types (**Figure 2D**). Termination of propagation at the seed nodes by the prize-collecting Steiner tree algorithm is the reason for recovering fewer driver genes in networks inferred by OI2. In large reference networks, highly connected nodes generate network shortcuts instead of using signal cascades or motifs. In PL-generated networks, when recruiting the shortest paths, intermediate nodes corresponded more to highly connected genes than specific driver genes. In pyPARAGON, we use the PageRank algorithm to propagate seed nodes to the neighbors in the reference interactome, which helps in obtaining more candidate drivers. Additionally, GGN construction filters out possible “frequent flyers” with the help of graphlets, which enable us to predict driver genes more precisely. Overall, pyPARAGON performs significantly better in cancer driver network prediction and can be further elaborated for tumor- or patient-specific network construction and network similarity-based comparisons.

### Patient-Specific Network Inference Unveil Hidden Commonalities across Tumors

We used pyPARAGON to construct the specific networks for 105 breast cancer patients’ tumors where the seed nodes are significantly altered phosphoproteins. The reference interactome is HIPPIE v2.3 For each tumor-specific network, we detected the modules -i.e. communities-with Louvain’s algorithm. We consider the modules as functional subunits of networks that individually or in cooperation take part in context-specific molecular processes. pyPARAGON uses hypergeometric tests to identify these active modules that are significantly over-represented in specific biological processes (**Supplementary Figure 2**). **Figure 3** shows an example patient-specific network composed of active modules that is organized based on the cellular localization of nodes. Modules have cross-talk between each other (inter-module edges). Many modules have proteins from different cellular compartments and are enriched in multiple biological processes. We calculated patient similarities using cosine similarity scores based on significant biological processes. This is a data reduction approach where we first translated a given list of proteins to a context-specific network, then divided it into modules and find the significant processes and finally these biological processes are used for similarity calculation across the tumors. t-SNE algorithm is used to represent tumor similarities based on two components. Eventually tumors are clustered into four groups (**Figure 4A**). **Supplementary Table 2** lists the 20 most frequently identified biological functions for each cluster. We uncovered significant biological processes that are present in at least two clusters (**Figure 4B**). In patient cluster-1, the most frequently associated biological process is the ubiquitin-dependent protein catabolic process, where several transcription factors and enzymes are present. Ubiquitination (one of the post-translational modifications) is a multistep enzymatic process involved in the regulation of cancer metabolism, such as the cell cycle, DNA damage repair, chromatin remodeling, and several signaling pathways ^51^. Ubiquitin-specific peptidases (USP) are regarded as potential therapeutic targets. However, any USP inhibitor is not in the clinical trial stage despite having promising potential in breast cancer ^52,53^. The patients in cluster-2 frequently share the mitotic cytokinesis process. Cytokinesis defects increase chromosomal instability, vast genomic alteration, and point mutations, and so provoke intratumoral heterogeneity ^54,55^. As shown in the patient similarity network (Supplementary Figure 3), only five patients have high similarity due to heterogeneity. Interestingly, we found that the nervous system development (NSD) process was the most frequent biological process in cluster-3. Breast cancer is the second most common cause of central nervous system (CNS) metastasis after lung cancer ^56^. In our datasets, just two patients had metastases. We found both patients with the NSD process in cluster-3. In cluster-4, we frequently found the regulation process of actin cytoskeleton organization, which is critically involved in cancer initiation, metastasis, and therapeutic responses. Rho GTPases, a family of the Ras GTPase superfamily, play a key role in this regulation ^57^. We also performed a Kaplan-Meier survival analysis across four patient clusters and did not find any significant differences (**Supplementary Figure 4**). However, in a pairwise comparison, we found that patients in cluster-4 have a significantly worse survival probability than cluster-1 (**Figure 4C**). Followingly, we identified active modules with KEGG pathway information to figure out over-represented pathways in these clusters (**Figure 4D**) ^58^. Cell cycle, oocyte meiosis, and PI3K/Akt signaling pathways are the most frequent and common pathways in clusters, except for cluster-2, while more frequent in cluster-1 than cluster-4. On the other hand, focal adhesion and Ras signaling pathways are significantly more frequent in cluster-4. The Ras signaling pathway is one of the key pathways for drug resistance owing to the bypassing of drug action mechanisms in the signaling network ^48,49^. In **Figure 4E**, we demonstrated the module associated with the Ras-signaling pathway, where pyPARAGON linked phosphoproteins with intermediate nodes including KRAS, NRAS, HRAS, RHOA, and RHOD. Next, we incorporated drugs targeting the active modules in the network. We collected drug-target interactions from the Therapeutic Target Database into modules ^50^. We extracted 8297 drugs, 330 drug targets, and active modules associated with 161 pathways for 105 breast cancer patients (**Supplementary Table 3**). An example of context-specific drugs for the active modules of patient A2-A0YD is shown in **Figure 5**. Adagrasib (MRTX849) and Sotorasib specifically target the Ras signaling-associated module. Both drugs are novel KRAS^G12C^ inhibitors approved by the FDA ^59,60^.

**Figure 3.**
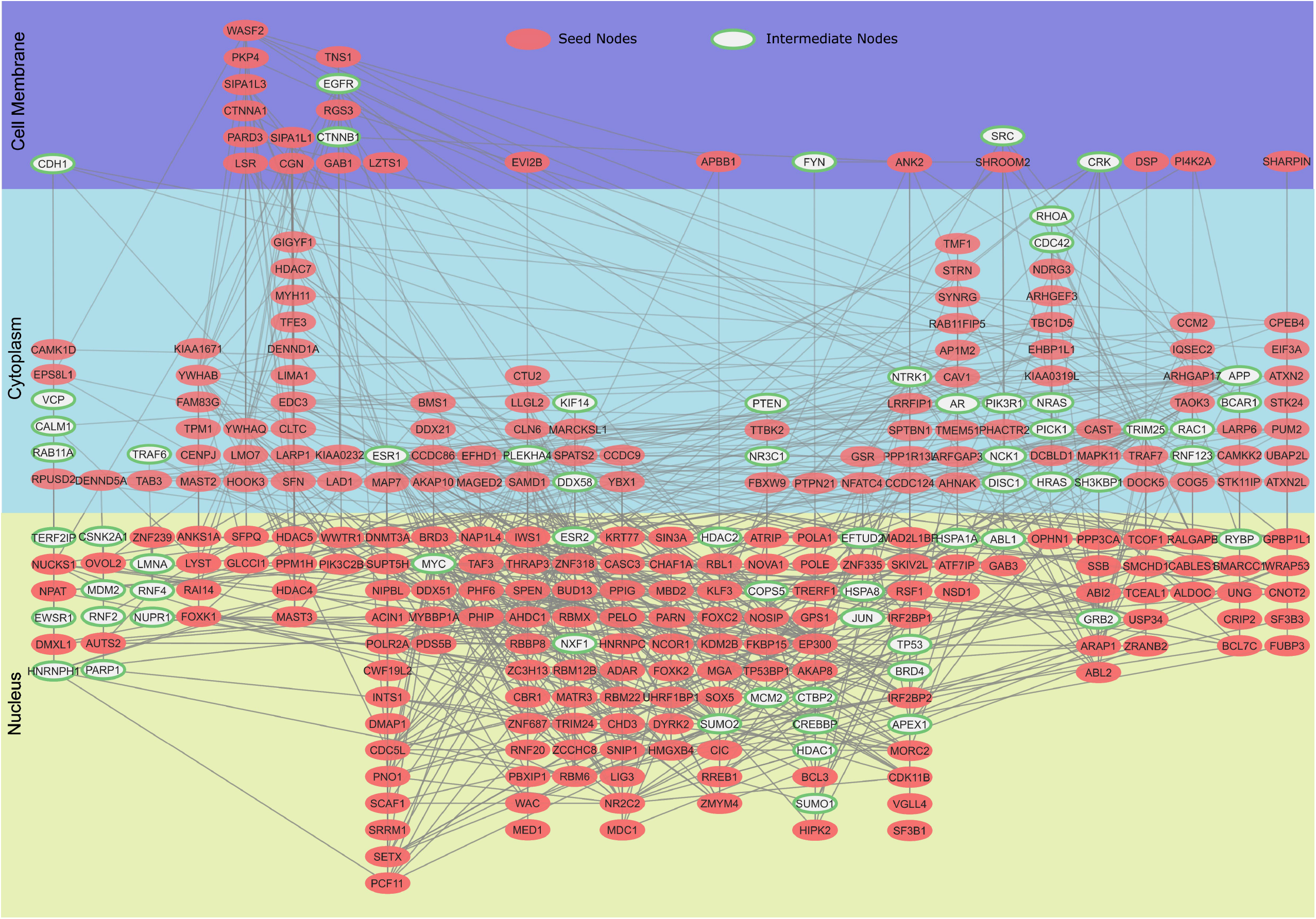
An example patient-specific network that is constructed by pyPARAGON (TCGA-A2-A0YD). Significantly phosphorylated proteins were used as the initial (seed) node set (colored red), and intermediate nodes predicted by pyPARAGON are green. Active modules that are associated with at least one significantly overrepresented biological process are shown in this patient-specific network. Each vertically aligned node set represents one active module. Nodes in each active module are layered based on their subcellular localization.

**Figure 4.**
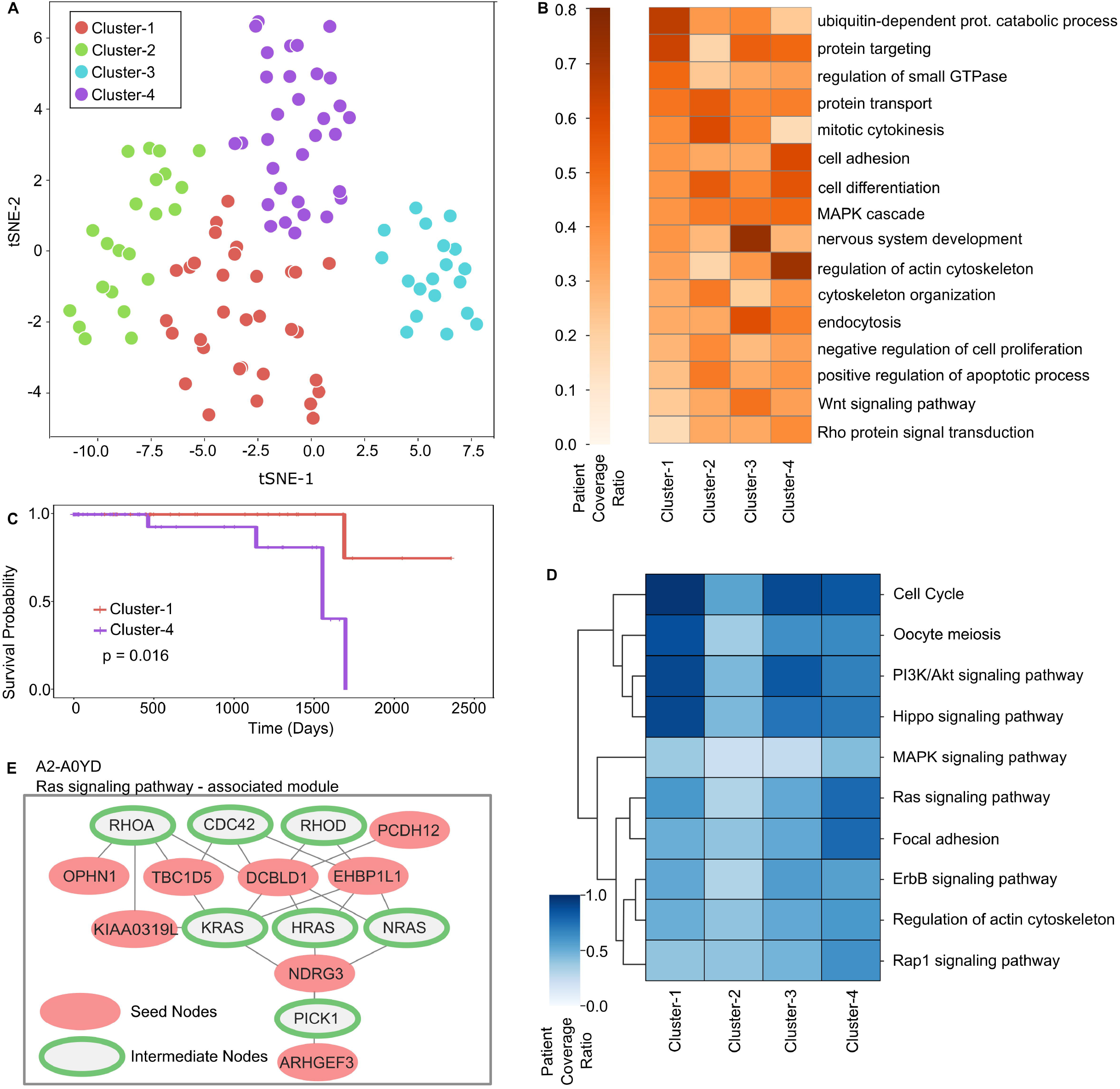
Stratification of tumors and associated biological processes with patient clusters. **a.** 105 breast cancer tumors are stratified into four clusters based on their similarity of significant biological processes in their network modules: cluster-1 (32 patients), cluster-2 (22 patients), cluster-3 (19 patients), and cluster-4 (32 patients). **b.** Heatmap of patient coverage ratio for each cluster and significant process pairs. A biological process is included in the heatmap if it is enriched in at least two clusters. The patient coverage ratio represents the ratio of patients having the enriched biological process in the corresponding clusters. The ubiquitin-dependent protein catabolic process and protein targeting were predominantly present in cluster-1 while mitotic cytokinesis in cluster-2, nervous system development in cluster-3, and actin cytoskeleton organization in cluster-4. **c.** Kaplan-Meier analysis shows the survival probabilities of cluster-1 (red) and cluster-4 (purple). Patients in cluster-4 have significantly worse survival in cluster-1. **d.** Heatmap shows significantly enriched KEGG pathways in active modules. The cell cycle, oocyte meiosis, PI3K/Akt, and Hippo signaling pathways were mostly observed in clusters-1, -3, and -4 while focal adhesion and Ras signaling pathways were prominent in cluster-4. **e.** The example module of A2-A0YD network corresponding to the Ras signaling pathway is shown where seed nodes are red and intermediate nodes are green.

**Figure 5.**
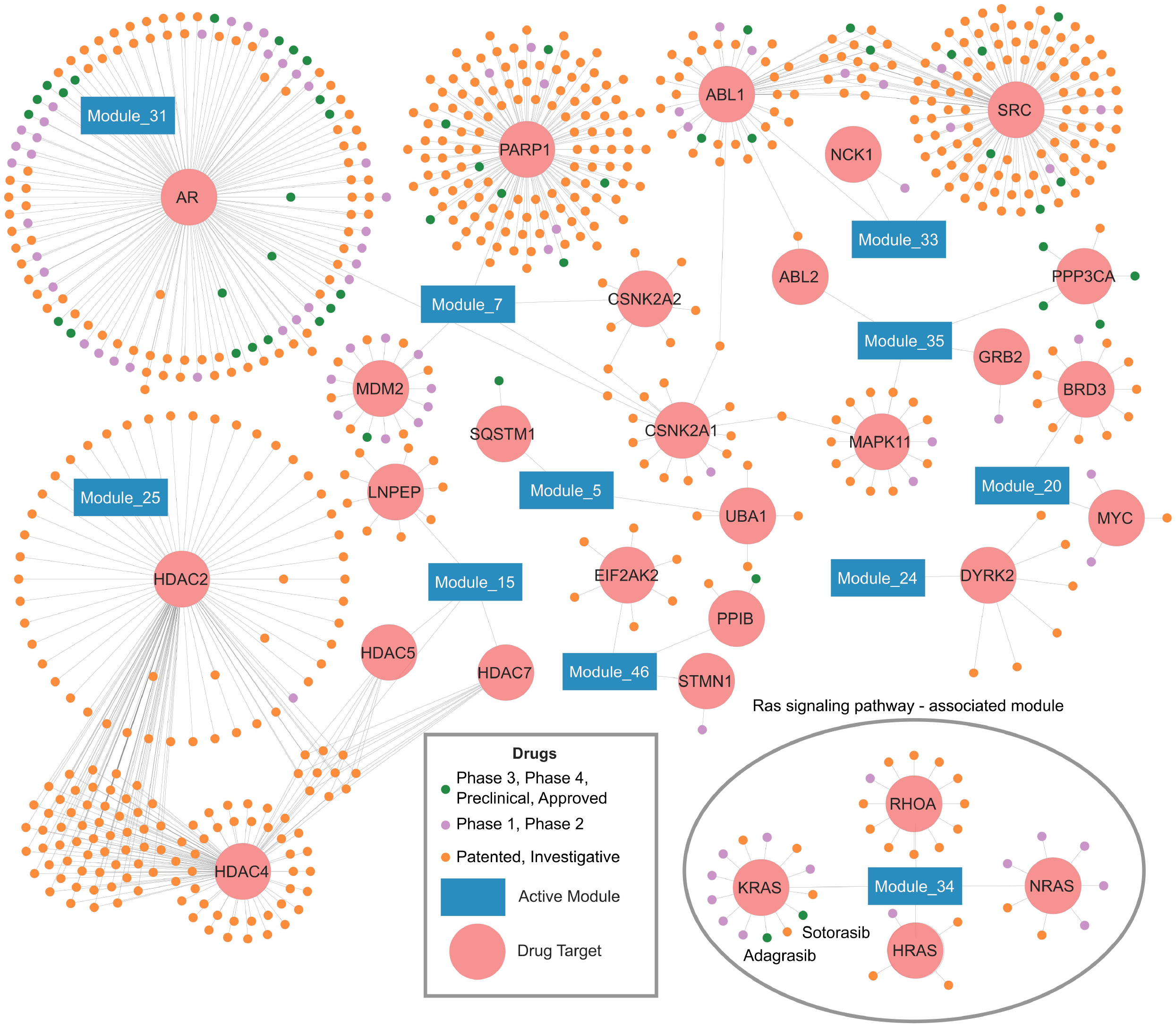
Drug-module interaction network of a patient (TCGA-A2-A9YD). Drugs are shown in three colors corresponding to three categories: drugs in phase 3, 4, or preclinical stage, and authorized drugs in green; drugs in phase 2 or 3 in purple and patented and investigational drugs are in pink. Drugs are connected to their immediate targets (light red circles) and active modules having these targets (blue rectangles) in the network. One example is the drugs Adagrasib (MRTX849) and Sotorasib which targets Module_34. This module is enriched in the Ras signaling pathway has the immediate target protein KRAS of these drugs.

## Discussion

In this work, we present pyPARAGON as a network-based multi-omic data integrator. pyPARAGON simultaneously utilizes the most frequent graphlets covering omic hits and network propagation to construct context-specific networks. Network inference algorithms encounter challenges arising from sparse data and the complexity associated with the growing number of interactions within reference networks. In our study, we address these issues by reducing reference networks to less complex networks composed of significantly frequent graphlets. By employing pyPARAGON, we mitigated the impact of noise generated by highly connected nodes in the reference networks. While reducing the noise, pyPARAGON preserves scale-free properties inherent in biological networks in the constructed GGNs. Additionally, we leveraged the PageRank flux calculation for edge prioritization and integrated GGNs with PageRank flux to successfully construct context-specific networks. By predicting driver genes, we extended the missing value problem in cancer-specific network construction. pyPARAGON inferred networks that encompassed a more precise and higher number of cancer drivers. Additionally, after inferring context-specific networks from phosphoproteomics, pyPARAGON can integrate modules and different types of annotations, such as biological processes, pathways, and drug knowledge. These findings indicate that pyPARAGON can predict cancer biomarkers, drivers, drugs, and therapeutic targets.

Different molecular aberrations, particularly in cancer, might result in identical disease manifestations ^61–63^. We used omics data of breast cancer tumors in CPTAC ^64^ as a case study to infer context-specific networks where interacting protein modules govern various biological processes and pathways. Patients were clustered based on the biological processes that were overrepresented in functional communities. We show that functional communities with the same driver genes mediate various biological processes according to the recruited proteins. Therefore, pyPARAGON is a powerful tool to identify disease-related molecular alterations and driver networks.

Despite the success of integrative approaches, including pyPARAGON, there are still issues in network-based omic data integration that must be addressed in the long-term. First, reference interactomes are incomplete ^65^. Notably, network-based methods strongly depend on features and coverage of reference networks ^66^. As a result of incomplete knowledge in large reference interactomes, protein complexes tend to form more topological modules than metabolic pathways ^67^. Thus, generic biological processes, such as transcription, replication, can be found more frequent in inferred networks. Thus, biological interpretations of context specific networks are challenging through causal relations, modules, and biological processes. Additionally, network-based methods do not assess the alternative copies of individual hits while various protein isoforms, post-translational modifications are within the proteome. Despite delivering more specific functions, this information is generalized and potentially lost in the network.

As a result of extended integrations in reference networks, missing interactions and highly connected nodes have become a prominent challenge in recent network inference tools based on belief propagation ^68,69^, random walks ^18,19,70^, the prize-collecting Steiner Forest ^17,71,72^, heat diffusion ^73,74^, and shortest path algorithms ^15,75^. Here, graphlets were deployed in our approaches for network trimming. In pathway reconstruction and the inference of context-specific networks, we compared our method with two popular tools: PL and OI2. Hub proteins may dominate the inferred network with unrelated interactions. The prize-collecting Steiner Forest algorithm penalizes hubs based on the number of interactions. Similarly, the flux calculation in pyPARAGON is a countermeasure against the curse of hubs beyond scoring interactions. OI2 and pyPARAGON work better at predicting interactions. In terms of the identification of associated genes, our tool outperformed the other tools. In the PL algorithm, highly connected nodes further diminish the shortest paths between seed nodes. OI2 early terminates the propagation of the seed nodes in a large reference network. However, the PageRank algorithm in pyPARAGON propagates the seed nodes before network inference, independent of GGN. Thus, pyPARAGON optimizes the inference of interactions and the propagation of seed nodes in the network.

Here, we only utilized graphlets composed of interactions among 3 and 4 nodes rather than interactions between 2 nodes. However, various graphlet information in reference networks, such as graphlet degree distribution, graphlet frequencies, and probabilistic graphlets, can be embedded in network inference algorithms or biological interpretations ^32,34,35,39,40^. However, the use of graphlet features will come with a high computational cost. To enhance context specificity, permutation-based methods can be additionally applied in downstream analysis where hypergeometric tests on communities are only used ^16^. The placement of community members in subcellular locations shows that inferred networks are composed of several signaling cascades from cell receptors to regulatory proteins. These communities can be detailed with mechanistic and causal relations for downstream analysis.

In conclusion, we released a novel tool, pyPARAGON, which infers context-specific networks by using graphlets and network propagation. It mitigates noise generated by highly connected nodes, preserves scale-free properties, and integrates network modules and biological annotations. Through its inferred network, we can predict context-specific biomarkers, drugs, and therapeutic targets. For downstream analysis, communities in the network can potentially be used to identify mechanistic molecular relations in complex and rare diseases. Here, pyPARAGAON integrated bulk omic data for static patient-specific network models. The next version of pyPARAGON will be an extension to integrate omic data at the single cell level to elucidate cell-type specific interactions.

## Methods

### Interactome and Datasets

We separately used interactomes as references; HIPPIE v2.2 (15 861 nodes, 345 770 edges), HIPPIE v2.3 (19 437 nodes, 774 449 edges), and ConsensusPathDB v35 (18 178 nodes, 516 211) ^76,77^. As a benchmark, we utilized 18 cancer signaling pathways in NetPath that are composed of more than 50 proteins ^47^. We prepared the seed node set for 8 cancer types; bladder urothelial carcinoma (BLCA), breast invasive carcinoma (BRCA), esophageal carcinoma (ESCA), head and neck squamous cell carcinoma (HNSC), lung squamous cell carcinoma (LUSC), pancreatic adenocarcinoma (PAAD), prostate adenocarcinoma (PRAD), and uterine corpus endometrial carcinoma (UCEC). We selected the 300 genes with the most frequent mutations among 1,289,655 mutations belonging to 3759 patients. The mutation dataset covers various cancer genomics projects, including TCGA and GENIE ^78^. We retrieved the 3333 driver mutations from IntOGen, harbored on 568 genes ^79^.

We retrieved phosphoproteomic data from 105 breast cancer patients and three healthy samples ^64^. We selected phosphosites that were at least identified in 50% of samples and had a standard deviation larger than 0.5 across all normalized samples. Phosphoproteins were categorized based on two criteria. (1) a higher log-2-fold-change (LFC) than 2, and (2) highly or less phosphorylated in the Gaussian Mixture Model (GMM) ^80^. In GMM, we split phosphoproteomics into three divisions: highly, less, or normally phosphorylated proteins. We ran the model 100 times at random for each phosphoproteomics and chose highly or less phosphorylated proteins in 95% of the models. Using the unit-variance scaling method on LFC, we gave differential phosphoproteins scores between 0.5 and 1 ^81^. We utilized biological processes retrieved from gene ontology, pathways from KEGG, subcellular localization from the human proteome atlas, and drug information from the therapeutic target database ^19,58,82,83^.

## Network Inference Methods

### PageRank-flux on Graphlet-Guided Network

We used 2-, 3-, and 4-node-graphlets (***G_0_, G_1_, G_2_, …, G_8_,*** shown in **Figure 1B**), which are small non-isomorphic subgraphs. An isomorphism of graphlets between two subgraphs, ***X(V_X_, E_X_)*** and ***Y(V_Y_, E_Y_)***, is defined with bijections between ***V_X_*** and ***V_Y_*** ^36^. We searched the graphlets for an intermediate node in one of the highest-degree orbits and seed nodes in the remaining orbits. The reference network is ***R*(*V_R_, E_R_, c(e)*),** where ***V_R_***, ***E_R_,*** and ***c(e)*** are nodes, undirected edges, and the confidence score of an edge, respectively. Similarly, we calculated the frequencies of graphlets in 100 permuted networks, recruiting the same seed node set. We compared the targeting graphlet frequencies in the reference and permuted networks with a z-test (p<0.05, z-score > 1.65). The union of graphlet motifs, a significantly high number of graphlets, constructs the graphlet-guided network (GGN), ***G(V_G_, E_G_),*** where ***G*** ⊆ ***R***.

The Personalized PageRank (PPR) algorithm calculates the probability of being at the node ***y***, ***p(y),*** at a particular time step (***t***), in the reference interactome according to formula 1, where the damping factor (λ) defines the possibility of walking from neighbor nodes (***x_i_***) to ***y***, and ***N*** is the number of nodes ^84,85^.

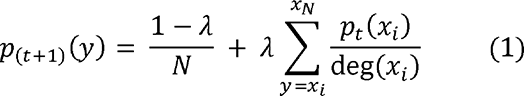

We calculated the directional flux scores for both directions (***f_u_***_→_***_t_*** and ***f_t_***_→_***_u_***) by using formulas 2 and 3, where ***u, t*** ∈ ***V_R_*** and ***e*** is the edge between ***u*** and ***t***, and ***deg(u)***, ***deg(t)*** are the number of neighbors of nodes, ***u*** and ***t***, respectively ^44^. The negative logarithm of minimum flux scores is used as a final edge (***f(e)***) score defined in formula 4.

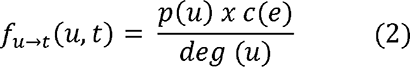

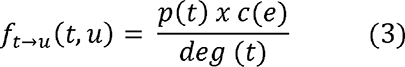

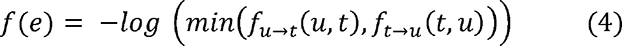

We weighted the edge set of GGN, ***G(V_G_, E_G_)***, with ***f(e)*** where ***e_1_, e_2_, e_3_, …, e_j_,…e_n_*** ∈ ***E_G_***, ***1*** ≤ ***j*** ≤ ***n*** and ***f(e_j-1_)> f(e_j_)>f(e_j+1_).*** The total flux scores (***F***) in GGN are calculated as formulated in formula 5.

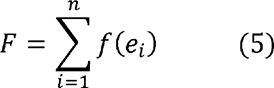

Let τ (***0*** ≤ τ ≤ ***1***) represent the scaling factor describing the threshold percentage of ***F***. We selected the edges by summing flux scores up to τ***xF*** (formula 6). In this way, we infer the context-specific network ***C(V_C_, E_C_),*** where ***E_C_***⊆ ***(E_G_,>)*** and ***V_C_***⊆ ***V_G_***.

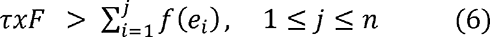

### Performance Assessment of Network Inference

We compare pyPARAGON with Omics Integrator 2 (OI2) and PathLinker 1.4.3 (PL) by reconstructing pathways in NetPath and inferring specific cancer networks.

OI2 implements the prize-collecting Steiner Forest algorithm ^86^. The objective function of OI2 combines confidence scores of edges ***(c(e)***) and penalties of edges calculated with node degrees and the scale parameter, □. The following function finds an optimum forest, ***F(V, E)***, by minimizing the objective function, the formula 7,

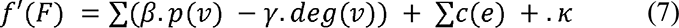

where *κ* is the number of connected components, *β* controls the relative weight of the node prizes, *μ* affects the penalty for the degree of a node represented by ***deg(v),*** and *ω* controls the cost of adding a tree to the solution network.

PL computes the ***k***-highest scoring short paths between seed nodes without a loop in the reference network. The path score, ***W***, is the product of the edge weights along the path ^15,87^. PL calculates the cost of a path with the formula 8:

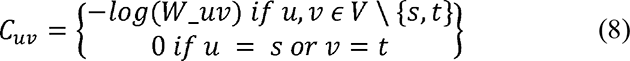

where ***s*** and ***t*** are, respectively, a source and a target for each node, x ∈ S. The cost of a path is the sum of the costs of the edges in the path.

Each pathway in NetPath was independently shuffled, and their nodes were split into two equal parts, five times. We reconstructed the pathways, recruiting one part as a seed nodeset and HIPPIE v2.2 as a reference.

The driver genes were randomly divided for each cancer type into five equal portions. Each portion was removed from the most frequently mutated genes. Then, we utilized the remaining frequently mutated genes as a seed nodeset and HIPPIE v2.2 as a reference so as to infer cancer specific networks.

We separately calculated precision, recall, F1 scores, and area under the precision-recall curve (AUPRC) for each pathway and each cancer specific network ^42^. We created grid search parameter sets for tools. In OI2, parameter sets were ranged as following; dummy edge weight *(ω),* edge reliability *(β)* between 0 and 5 with 0.5 increments, and degree penalty *(□)* between 0 and 10 with increment 1. Similarly, we measured the performance of PL by altering ***k***, the number of shortest paths, between 50-1000 with increments of 50, while the performance of pyPARAGON by ranging the damping factor (λ) and flux threshold (τ) between 0.05 and 1 with 0.05 increments.

To evaluate the performance of GGN, we quantified the alteration of highly connected proteins between the given reference network and GGN. We defined the highly connected proteins ***H_R_*** with more than 200 interactions, (***h_1_, h_2_, …, h_n_***) ∈ ***H_R_*** for a reference network, the highly connected proteins (***h_1_, h_2_, …, h_m_***) ∈ ***H_G_***in GNN, and the highly connected proteins (***h_1_, h_2_, …, h_p_***) ∈ ***H_P_***in the given pathway, ***H_P_*** ⊆ ***H_G_***⊆ ***H_R_*** ⊆ ***V_R_***. The reduction ratio (***RR***) of the remaining highly connected proteins in GGN was separately calculated using the formula 9:

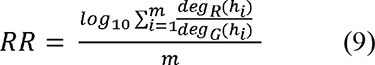

where ***deg(h)*** is the number of interactions of ***h*** and *m* is the number of highly connected nodes in GGN. We separately calculated the reduction ratio of highly connected proteins for each signaling pathway.

### Patient-Specific Network Construction

pyPARAGON constructed the patient-specific networks for 105 BRCA patients by recruiting differential phosphoproteins as seed nodes and HIPPIE v2.3 as a reference network ^64^. In the construction of GGN, we directly parsed graphlets that had been determined in pathway reconstruction.

We ran the Louvain method, a fast and heuristic method composed of two iterative steps. (1) Assigning each node to its community; and (2) Interchanging neighbor nodes to find the maximum modularity until no positive gain is achieved ^85^.

We investigated the over-represented biological processes, KEGG pathways in the inferred networks. We utilized the hypergeometric distribution, which describes the probabilities of communities associated with the target pool, such as pathways or biological processes. We calculated the p-value using the formula 10 ^46^.

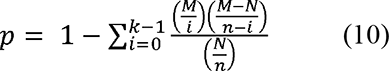

We define M as the population size, the number of genes in the reference network; ***N*** as the number of genes in the target pool; ***n*** as the number of genes in the community; and ***k*** as the number of successfully identified genes in the target process. We only selected the most significant community for each biological process, or KEGG pathway since multiple communities pointed out the same process. Then, we eliminated insignificant communities and their associated biological processes and pathways.

For each patient, we constituted the vector of biological processes, which was only composed of significant biological processes. We computed a similarity matrix that measured the pairwise cosine similarities between all paired patients. We applied the t-distributed stochastic neighbor embedding (t-SNE) algorithm to transform the similarity matrix into two-dimensional data, component-1, and component-2 ^88^. The patient groups were determined with agglomerative clustering through the euclidean distance.

We used "survminer", an R library, for the Kaplan-Meier survival curve, indicating the percentage of alive patients in the group over time ^89^. The log-rank test computes the chi-square (χ^2^) for each group at each event time and gathers their outputs in the result table. The final chi-square score and p-value are calculated by comparing the curves of each group. The log-rank p-value implies the ability of the model to differentiate two risk groups.

## Declarations

### Ethics approval and consent to participate

Not applicable.

### Consent for publication

Not applicable.

### Data availability

The results shown here are in part based upon data generated by the TCGA Research Network: https://www.cancer.gov/tcga. Open-source datasets were utilized in this paper. As reference networks, we retrieved HIPPIE v2.2 and v2.3 from http://cbdm-01.zdv.uni-mainz.de/~mschaefer/hippie/, ConsensusPathDB v35 from http://cpdb.molgen.mpg.de/. The signaling pathways were downloaded from NetPath, http://www.netpath.org/. Mutation frequencies for eight cancer types were available in ^75^. Cancer driver genes were retrieved from IntOGen, https://www.intogen.org. Phosphoproteomic data from 105 breast cancer patients was generated by Berliner ^56^. Biological process and pathway information were downloaded from http://geneontology.org/ and https://www.genome.jp/kegg/pathway.html, respectively. Drug information were retrieved from the therapeutic target database, https://db.idrblab.net/ttd/

### Competing interests

The authors declare that they have no competing interests.

## Acknowledgments

NT has received support from the 2247-A National Outstanding Researchers Program of TUBITAK under the project number 121C292. MKA has been financially supported by TUBITAK-2211 fellowship.

## Authors’ contributions

Conceptualization: MKA, NT

Data curation: MKA

Methodology: MKA, NT

Project administration: NT

Supervision: NT

Visualization: MKA, NT

## Notes

### Competing Interest Statement

The authors have declared no competing interest.

## References

1. Liu, A. et al. From expression footprints to causal pathways: contextualizing large signaling networks with CARNIVAL. NPJ Syst Biol Appl 5, 40 (2019).

2. Ross, K. E. et al. iPTMnet: Integrative Bioinformatics for Studying PTM Networks. Methods Mol. Biol. 1558, 333–353 (2017).

3. Heumos, L. et al. Best practices for single-cell analysis across modalities. Nat. Rev. Genet. 1–23 (2023).

4. Boehm, K. M., Khosravi, P., Vanguri, R., Gao, J. & Shah, S. P. Harnessing multimodal data integration to advance precision oncology. Nat. Rev. Cancer 22, 114–126 (2022).

5. Rautenstrauch, P., Vlot, A. H. C., Saran, S. & Ohler, U. Intricacies of single-cell multiomics data integration. Trends Genet. 38, 128–139 (2022).

6. Aalto, A., Viitasaari, L., Ilmonen, P., Mombaerts, L. & Gonçalves, J. Gene regulatory network inference from sparsely sampled noisy data. Nat. Commun. 11, 3493 (2020).

7. Ren, M., Pokrovsky, A., Yang, B. & Urtasun, R. SBNet: Sparse Blocks Network for Fast Inference. in Proceedings of the IEEE Conference on Computer Vision and Pattern Recognition, 8711–8720 (2018).

8. Demirel, H. C., Arici, M. K. & Tuncbag, N. Computational approaches leveraging integrated connections of multi-omic data toward clinical applications. Mol Omics 18, 7–18 (2022).

9. Tong, L., Mitchel, J., Chatlin, K. & Wang, M. D. Deep learning based feature-level integration of multi-omics data for breast cancer patients survival analysis. BMC Med. Inform. Decis. Mak. 20, 225 (2020).

10. Kim, S. Y., Jeong, H.-H., Kim, J., Moon, J.-H. & Sohn, K.-A. Robust pathway-based multi-omics data integration using directed random walks for survival prediction in multiple cancer studies. Biol. Direct 14, 8 (2019).

11. Duan, R. et al. Evaluation and comparison of multi-omics data integration methods for cancer subtyping. PLoS Comput. Biol. 17, e1009224 (2021).

12. Malod-Dognin, N. et al. Towards a data-integrated cell. Nat. Commun. 10, (2019).

13. Shah, H. A., Liu, J., Yang, Z., Zhang, X. & Feng, J. DeepRF: A deep learning method for predicting metabolic pathways in organisms based on annotated genomes. Comput. Biol. Med. 147, 105756 (2022).

14. Costello, Z. & Martin, H. G. A machine learning approach to predict metabolic pathway dynamics from time-series multiomics data. NPJ Syst Biol Appl 4, 19 (2018).

15. Ritz, A. et al. Pathways on demand: Automated reconstruction of human signaling networks. npj Systems Biology and Applications 2, 1–9 (2016).

16. Levi, H., Elkon, R. & Shamir, R. DOMINO: a network-based active module identification algorithm with reduced rate of false calls. Mol. Syst. Biol. 17, e9593 (2021).

17. Tuncbag, N. et al. Network-Based Interpretation of Diverse High-Throughput Datasets through the Omics Integrator Software Package. PLoS Comput. Biol. 12, (2016).

18. Jagtap, S. et al. BraneMF: integration of biological networks for functional analysis of proteins. Bioinformatics 38, 5383–5389 (2022).

19. Zhou, Y. et al. Therapeutic target database update 2022: facilitating drug discovery with enriched comparative data of targeted agents. Nucleic Acids Res. 50, D1398–D1407 (2022).

20. 20. Cowen, L., Ideker, T., Raphael, B. J. & Sharan, R. Network propagation: A universal amplifier of genetic associations. Nature Reviews Genetics vol. 18, 551–562 Preprint at https://doi.org/10.1038/nrg.2017.38 (2017).

21. Di Nanni, N., Bersanelli, M., Milanesi, L. & Mosca, E. Network Diffusion Promotes the Integrative Analysis of Multiple Omics. Frontiers in Genetics 10.3389/fgene.2020.00106 (2020).

22. Reyna, M. A., Leiserson, M. D. M. & Raphael, B. J. Hierarchical HotNet: identifying hierarchies of altered subnetworks. Bioinformatics 34, i972–i980 (2018).

23. Silverman, E. K. et al. Molecular networks in Network Medicine: Development and applications. Wiley Interdisciplinary Reviews: Systems Biology and Medicine vol. 12 1489 Preprint at https://doi.org/10.1002/wsbm.1489 (2020).

24. Luna, A. et al. Analyzing causal relationships in proteomic profiles using CausalPath. STAR Protoc 2, 100955 (2021).

25. Dugourd, A. et al. Causal integration of multiComics data with prior knowledge to generate mechanistic hypotheses. Mol. Syst. Biol. 17, e9730 (2021).

26. Levitsky, L. I. et al. IdentiPy: An Extensible Search Engine for Protein Identification in Shotgun Proteomics. J. Proteome Res. 17, 2249–2255 (2018).

27. Unsal-Beyge, S. & Tuncbag, N. Functional stratification of cancer drugs through integrated network similarity. NPJ Syst Biol Appl 8, 11 (2022).

28. Nussinov, R. et al. Mechanism of activation and the rewired network: New drug design concepts. Med. Res. Rev. 42, 770–799 (2022).

29. Dincer, C., Kaya, T., Keskin, O., Gursoy, A. & Tuncbag, N. 3D spatial organization and network-guided comparison of mutation profiles in Glioblastoma reveals similarities across patients. PLoS Comput. Biol. 15, e1006789 (2019).

30. Hristov, B. H., Chazelle, B. & Singh, M. uKIN Combines New and Prior Information with Guided Network Propagation to Accurately Identify Disease Genes. Cell Systems 10, 470–479.e3 (2020).

31. Ogris, C., Hu, Y., Arloth, J. & Müller, N. S. Versatile knowledge guided network inference method for prioritizing key regulatory factors in multi-omics data. Sci. Rep. 11, 6806 (2021).

32. Yaveroğlu, Ö. N. et al. Revealing the hidden language of complex networks. Sci. Rep. 4, 4547 (2014).

33. Wong, S. W. H., Cercone, N. & Jurisica, I. Comparative network analysis via differential graphlet communities. Proteomics 15, 608–617 (2015).

34. Sarajlić, A., Malod-Dognin, N., Yaveroğlu, Ö. N. & Pržulj, N. Graphlet-based Characterization of Directed Networks. Sci. Rep. 6, 35098 (2016).

35. Martin, A. J. M., Dominguez, C., Contreras-Riquelme, S., Holmes, D. S. & Perez-Acle, T. Graphlet Based Metrics for the Comparison of Gene Regulatory Networks. PLoS One 11, e0163497 (2016).

36. Przulj, N. Biological network comparison using graphlet degree distribution. Bioinformatics 23, e177–83 (2007).

37. Windels, S. F. L., Malod-Dognin, N. & Pržulj, N. Graphlet Laplacians for topology-function and topology-disease relationships. Bioinformatics 35, 5226–5234 (2019).

38. Li, Q. & Milenkovic, T. Supervised Prediction of Aging-Related Genes From a Context-Specific Protein Interaction Subnetwork. IEEE/ACM Trans. Comput. Biol. Bioinform. 19, 2484–2498 (2022).

39. Zhang, L., Liu, T., Chen, H., Zhao, Q. & Liu, H. Predicting lncRNA-miRNA interactions based on interactome network and graphlet interaction. Genomics 113, 874–880 (2021).

40. Magnano, C. S. & Gitter, A. Automating parameter selection to avoid implausible biological pathway models. npj Syst Biol Appl 7, 12 (2021)

41. Babur, Ö. et al. Causal interactions from proteomic profiles: Molecular data meets pathway knowledge. Patterns 2, (2021).

42. Arici, M. K. & Tuncbag, N. Performance Assessment of the Network Reconstruction Approaches on Various Interactomes. Front. Mol. Biosci. 8, 666705 (2021).

43. Huang, J. K. et al. Systematic Evaluation of Molecular Networks for Discovery of Disease Genes. Cell Syst 6, 484–495.e5 (2018).

44. Rubel, T. & Ritz, A. Augmenting signaling pathway reconstructions. in Proceedings of the 11th ACM International Conference on Bioinformatics, Computational Biology and Health Informatics 1, 1–10 (2020).

45. Blondel, V. D., Guillaume, J.-L., Lambiotte, R. & Lefebvre, E. Fast unfolding of communities in large networks. J. Stat. Mech. 2008, P10008 (2008).

46. Boyle, E. I. et al. GO::TermFinder--open source software for accessing Gene Ontology information and finding significantly enriched Gene Ontology terms associated with a list of genes. Bioinformatics 20, 3710–3715 (2004).

47. Kandasamy, K. et al. NetPath: A public resource of curated signal transduction pathways. Genome Biol. 11, (2010).

48. Hu, G., Wu, Z., Uversky, V. N. & Kurgan, L. Functional Analysis of Human Hub Proteins and Their Interactors Involved in the Intrinsic Disorder-Enriched Interactions. Int. J. Mol. Sci. 18, (2017).

49. Vidal, M., Cusick, M. E. & Szló Barabá Si, A.-L. Leading Edge Review Interactome Networks and Human Disease. Cell 144, 986–998 (2011).

50. Charmpi, K., Chokkalingam, M., Johnen, R. & Beyer, A. Optimizing network propagation for multi-omics data integration. PLoS Comput. Biol. 17, e1009161 (2021).

51. Cruz, L., Soares, P. & Correia, M. Ubiquitin-Specific Proteases: Players in Cancer Cellular Processes. Pharmaceuticals 14, (2021).

52. Shi, D. & Grossman, S. R. Ubiquitin becomes ubiquitous in cancer: emerging roles of ubiquitin ligases and deubiquitinases in tumorigenesis and as therapeutic targets. Cancer Biol. Ther. 10, 737–747 (2010).

53. Huang, M.-L., Shen, G.-T. & Li, N.-L. Emerging potential of ubiquitin-specific proteases and ubiquitin-specific proteases inhibitors in breast cancer treatment. World J Clin Cases 10, 11690–11701 (2022).

54. Lens, S. M. A. & Medema, R. H. Cytokinesis defects and cancer. Nat. Rev. Cancer 19, 32–45 (2019).

55. Li, J., Dallmayer, M., Kirchner, T., Musa, J. & Grünewald, T. G. P. PRC1: Linking Cytokinesis, Chromosomal Instability, and Cancer Evolution. Trends Cancer Res. 4, 59–73 (2018).

56. Ben-Zion Berliner, M., et al. Central nervous system metastases in breast cancer: the impact of age on patterns of development and outcome. Breast Cancer Res. Treat. 185, 423–432 (2021).

57. Haga, R. B. & Ridley, A. J. Rho GTPases: Regulation and roles in cancer cell biology. Small GTPases 7, 207–221 (2016).

58. Kanehisa, M., Furumichi, M., Sato, Y., Kawashima, M. & Ishiguro-Watanabe, M. KEGG for taxonomy-based analysis of pathways and genomes. Nucleic Acids Res. 51, D587–D592 (2023).

59. Hallin, J. et al. The KRAS Inhibitor MRTX849 Provides Insight toward Therapeutic Susceptibility of KRAS-Mutant Cancers in Mouse Models and Patients. Cancer Discov. 10, 54–71 (2020).

60. Zhang, S. S. & Nagasaka, M. Spotlight on Sotorasib (AMG 510) for Positive Non-Small Cell Lung Cancer. Lung Cancer 12, 115–122 (2021).

61. Peng, J., Zhou, Y. & Wang, K. Multiplex gene and phenotype network to characterize shared genetic pathways of epilepsy and autism. Sci. Rep. 11, 952 (2021).

62. Riller, Q. & Rieux-Laucat, F. RASopathies: From germline mutations to somatic and multigenic diseases. Biomed. J. 44, 422–432 (2021).

63. Muñoz-Maldonado, C., Zimmer, Y. & Medová, M. A Comparative Analysis of Individual RAS Mutations in Cancer Biology. Front. Oncol. 9, 1088 (2019).

64. Mertins, P. et al. Proteogenomics connects somatic mutations to signalling in breast cancer. Nature 534, 55–62 (2016).

65. Cheng, F. et al. Comprehensive characterization of protein-protein interactions perturbed by disease mutations. Nat. Genet. 53, 342–353 (2021).

66. Kang, Y. et al. HN-PPISP: a hybrid network based on MLP-Mixer for protein-protein interaction site prediction. Brief. Bioinform. 24, (2023).

67. Mosca, E. et al. Characterization and comparison of gene-centered human interactomes. Brief. Bioinform. 22, (2021).

68. Kirkley, A., Cantwell, G. T. & Newman, M. E. J. Belief propagation for networks with loops. Sci Adv 7, (2021).

69. Korkut, A. et al. Perturbation biology nominates upstream-downstream drug combinations in RAF inhibitor resistant melanoma cells. Elife 4, (2015).

70. Ietswaart, R., Gyori, B. M., Bachman, J. A., Sorger, P. K. & Churchman, L. S. GeneWalk identifies relevant gene functions for a biological context using network representation learning. Genome Biol. 22, 55 (2021).

71. Sychev, Z. E. et al. Integrated systems biology analysis of KSHV latent infection reveals viral induction and reliance on peroxisome mediated lipid metabolism. PLoS Pathog. 13, e1006256 (2017).

72. Dinstag, G. & Shamir, R. PRODIGY: personalized prioritization of driver genes. Bioinformatics 36, 1831–1839 (2020).

73. Kuenzi, B. M. & Ideker, T. A census of pathway maps in cancer systems biology. Nat. Rev. Cancer 20, 233–246 (2020).

74. Leiserson, M. D. M. et al. Pan-cancer network analysis identifies combinations of rare somatic mutations across pathways and protein complexes. Nat. Genet. 47, 106–114 (2015).

75. Licata, L. et al. SIGNOR 2.0, the SIGnaling Network Open Resource 2.0: 2019 update. Nucleic Acids Res. 48, D504–D510 (2020).

76. Alanis-Lobato, G., Andrade-Navarro, M. A. & Schaefer, M. H. HIPPIE v2.0: enhancing meaningfulness and reliability of protein-protein interaction networks. Nucleic Acids Res. 45, (2017).

77. Kamburov, A. & Herwig, R. ConsensusPathDB 2022: molecular interactions update as a resource for network biology. Nucleic Acids Res. 50, D587–D595 (2022).

78. Ellrott, K. et al. Scalable Open Science Approach for Mutation Calling of Tumor Exomes Using Multiple Genomic Pipelines. Cell Syst 6, 271–281.e7 (2018).

79. Martamp, F. et al. A compendium of mutational cancer driver genes. Nat. Rev. Cancer 20, 555–572 (2020).

80. Scrucca, L., Fop, M., Murphy, T. B. & Raftery, A. E. mclust 5: Clustering, Classification and Density Estimation Using Gaussian Finite Mixture Models. R J. 8, 289–317 (2016).

81. Raschka, S., Liu, Y., Mirjalili, V. & Dzhulgakov, D. Machine Learning with PyTorch and Scikit-Learn: Develop machine learning and deep learning models with Python. (Packt Publishing Ltd, 2022).

82. Carbon, S. & Mungall, C. Gene Ontology Data Reference. Zenodo (2018).

83. Chapple, C. E. et al. Extreme multifunctional proteins identified from a human protein interaction network. Nat. Commun. 6, 7412 (2015).

84. Langville, A. N. & Meyer, C. D. A Survey of Eigenvector Methods for Web Information Retrieval. Society for Industrial and Applied Mathematics 47, 135–161 (2005).

85. Page, L., Brin. S., Motwani, R. & Winograd T. The PageRank Citation Ranking: Bringing Order to the Web. (1998).

86. Tuncbag, N. et al. Simultaneous reconstruction of multiple signaling pathways via the prize-collecting steiner forest problem. Journal of Computational Biology 20, 124–136 (2013).

87. Gil, D. P., Law, J. N. & Murali, T. M. The PathLinker app: Connect the dots in protein interaction networks. F1000Res. 6, 58 (2017).

88. Cieslak, M. C., Castelfranco, A. M., Roncalli, V., Lenz, P. H. & Hartline, D. K. t-Distributed Stochastic Neighbor Embedding (t-SNE): A tool for eco-physiological transcriptomic analysis. Mar. Genomics 51, 100723 (2020).

89. Scrucca, L., Santucci, A. & Aversa, F. Competing risk analysis using R: an easy guide for clinicians. Bone Marrow Transplant. 40, 381–387 (2007).

