## Supplementary Figures for "Unveiling Hidden Connections in Omics Data *via* pyPARAGON: an Integrative Hybrid Approach for Disease Network Construction"

### Supplementary Material

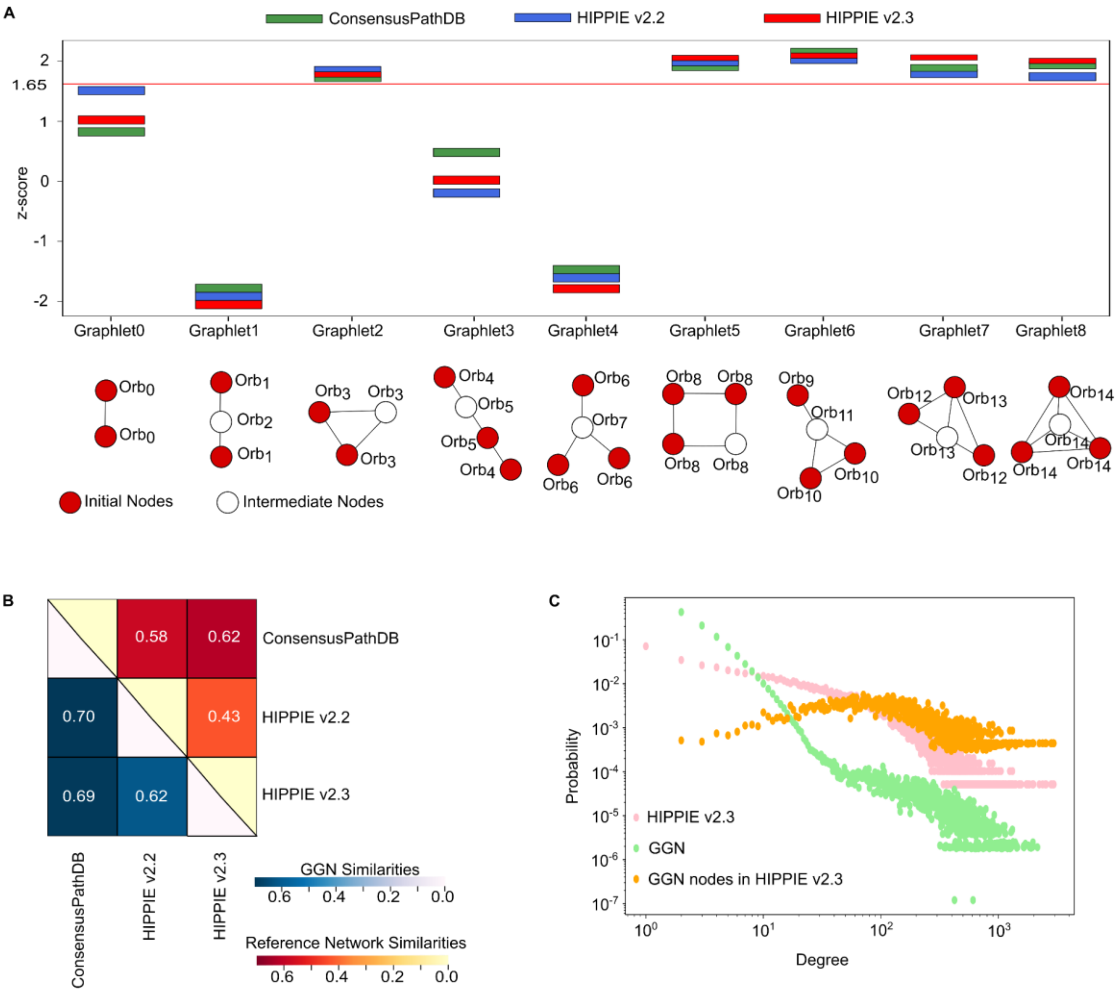


**Figure S1:** Graphlet-guided networks (GGN) optimize reference networks. **a.** Graphlets composed of 2, 3, and 4 nodes are constructed with initial nodes (red circle) coming from the given input and intermediate nodes (white circle). We compared the frequencies of graphlets on different reference interactomes with their 100 permuted networks. Despite having different network sizes and properties, ConsensusPathDB (green), HIPPIE v2.2 (blue), and HIPPIE v2.3 (red) have similar graphlet motifs, such as Graphlets 2, 5, 6, 7, and 8 for signaling pathways in NetPath ($p<0.05$). **b.** The heatmap with the gradual color change highlighted the network similarities between reference networks (red) and GGNs (blue). The Jaccard Similarity Index was determined by dividing the number of common interactions by the number of merged interactions. The top-right section depicts network similarities, whereas the bottom-left section depicts average GGN similarities retrieved with the same initial nodes from NetPath. The construction of GGN results in more comparable and optimized networks. **c.** Pink, green, and orange distributions depict the degree probabilities of HIPPIE v2.3, GGN, and GGN nodes in HIPPIE v2.3. A power law governs HIPPIE v2.3, a scale-free network. GGN was constructed with nodes at various degrees to avoid the noise of highly connected nodes by reducing their irrelevant interactions. GGN retains scale-free network features, as seen in biological networks.

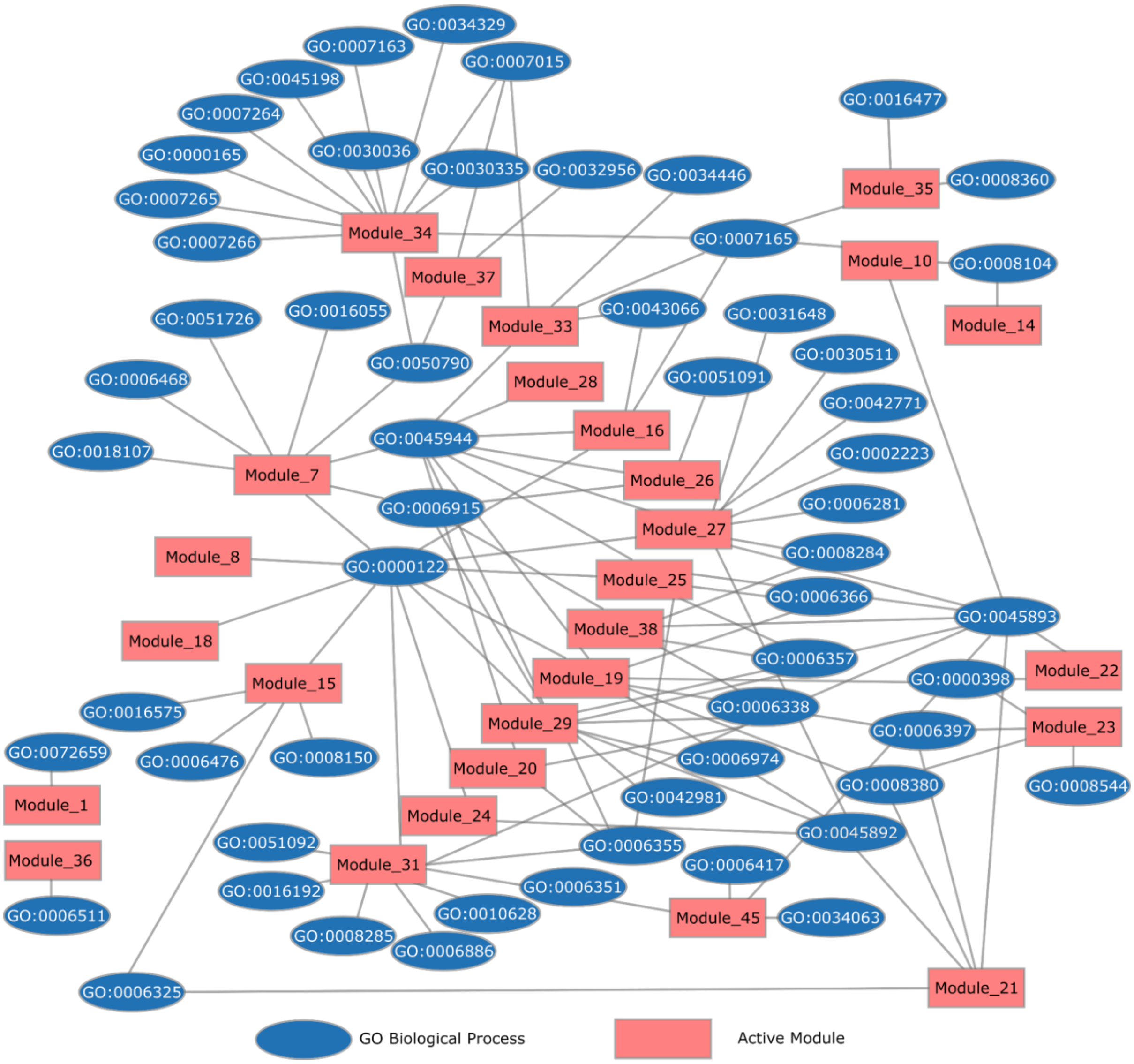


**Figure S2:** The association of active modules (pale red) and biological processes (blue) was illustrated in the network. An active module can be associated with multiple biological processes, or a biological process can be related to multiple active modules.

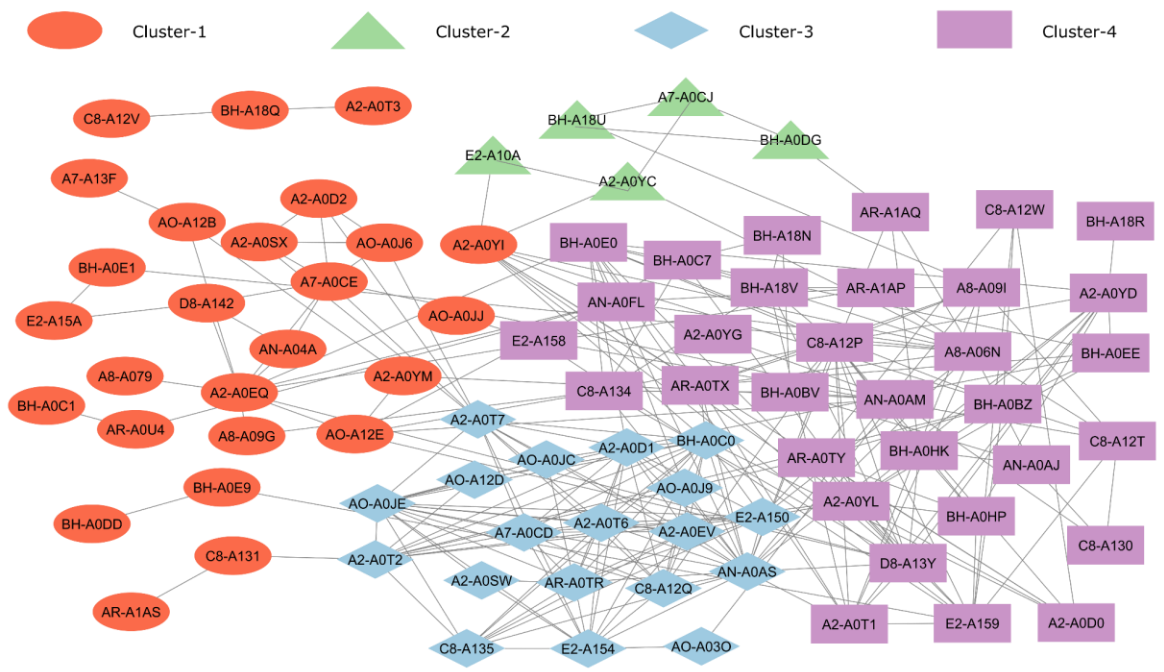


**Figure S3: Patient similarity network.** The similarities of 105 patients were calculated through a cosine similarity score of meaningful biological processes between patient pairs. In the similarity network, we illustrated interactions between patients with similarity scores greater than 0.5 (82 patients, 262 interactions). In the similarity network, we displayed interactions between patients with similarity scores greater than 0.5. (82 patients, 262 interactions). Using the t-SNE algorithm and agglomerative clustering, we divided the patients into four groups based on biological processes. Cluster-1 (26 patients) was represented by red ellipses, Cluster-2 (5 patients) by green triangles, Cluster-3 (19 patients) by blue diamonds, and Cluster-4 (32 patients) by purple rectangles. The majority of the patients in Cluster-2 do not have obvious similarities in patient pairs, while most in Cluster-3 and Cluster-4 do have higher similarities and more interactions in the patient similarity network.

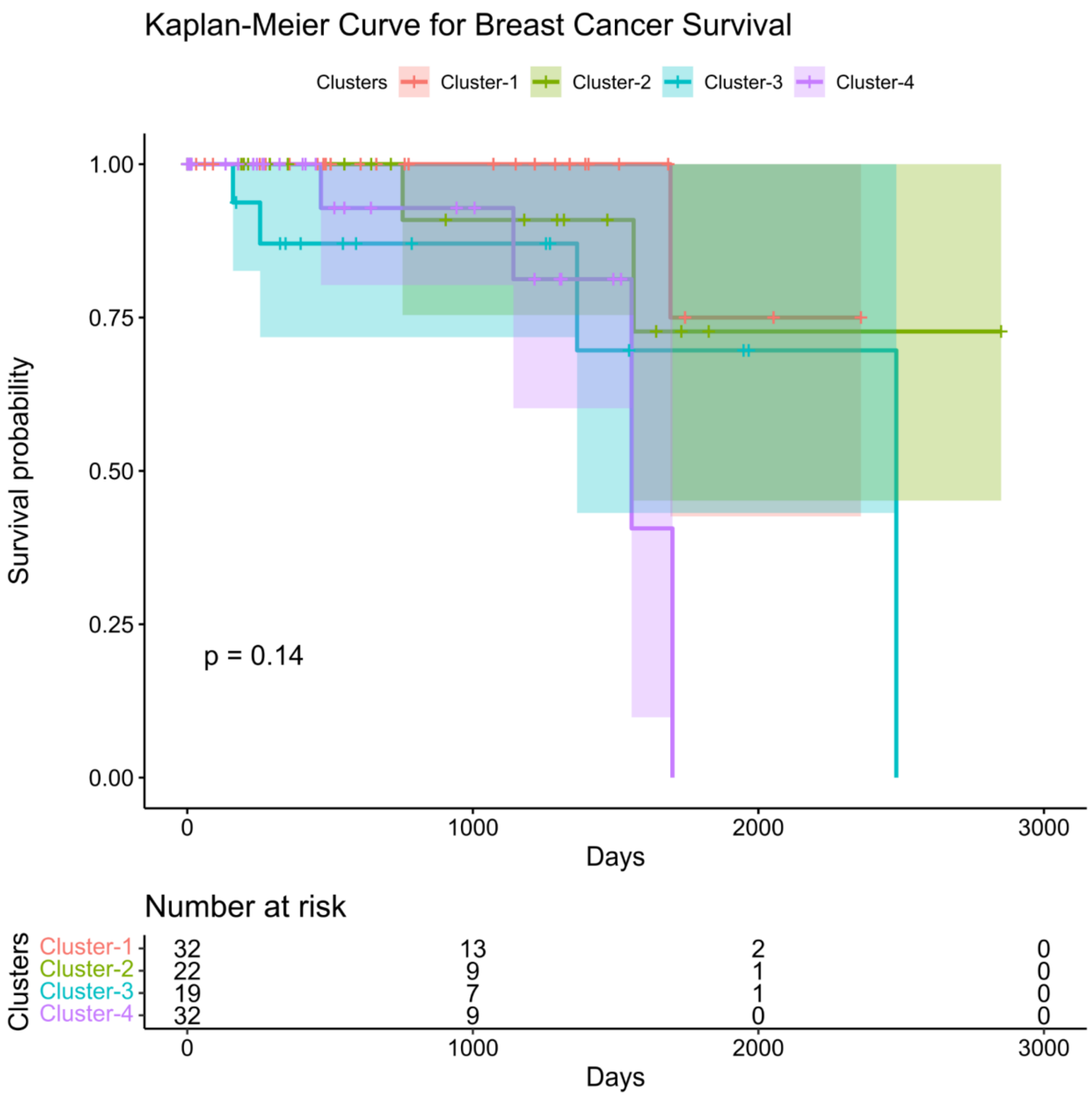


**Figure S4: The survival probabilities of clusters (A)** The Kaplan-Meier (K-M) curve analysis of clustered breast cancer showed that there was not any major difference in survival probabilities or risks among the four clusters. However, the lowest survival probability seems to be in cluster-4; the highest survival probability is in cluster-1.
